# Discovering 25 novel phyla that fill gaps in the eukaryotic tree of life

**DOI:** 10.64898/2026.08.28.747736

**Authors:** Leho Tedersoo, Vladimir Mikryukov, Sirje Sildever, Dominika Chmolowska, Kasia Piwosz, Mathisse Meyneng, Arthur Monjot, Javier del Campo, Enrique Lara, Ali Hakimzadeh, Stefan Geisen, Kristel Panksep, Mo Bahram, Angela M. Oliverio, Rachel M. Shepherd, Sonja Rückert, Anders Lanzen, Vedprakash G. Hurdeal, Maria Concetta Eliso, Raffaella Casotti, Mahdiehalsadat Hosseynimoghadam, Raffaele Siano, Marina Chauvet, Victoria Prins, Veljo Kisand, Sten Anslan, Saad Alkahtani, R. Henrik Nilsson

## Abstract

Protists play important roles in food chains and symbioses in soil and aquatic environments, displaying an enormous morphological and functional diversity^1,2^. While most commonly found protist species are well known to science, our global-scale environmental DNA survey across soil, water, and sediments reveals dozens of novel, phylum-level phylogenetic lineages that remain to be characterized for basic morphology and function. A vast majority of these undescribed taxa occur in marine water and sediments, but some are common in soil. Most of these novel taxa have distinct substrate and habitat preferences and biogeographic patterns. To accord these lineages scientific agency and enable unambiguous scientific communication, we propose formal names for 150 species to phylum-level taxa from 25 deep lineages based on eDNA and rRNA gene long-read sequence information.

## Introduction

Eukaryotes include diverse micro- and macroorganisms, with body sizes ranging from <1 micron to hundreds of meters. Unicellular eukaryote lineages - protists - are phylogenetically more diverse than macroscopic groups, such as animals, plants, dikarya fungi and brown algae^1^. Protists are key contributors to global ecosystems across oceans, freshwater, soils and host-associated niches, with diverse functional roles as primary producers, consumers, decomposers, symbionts and pathogens^2^. Microscopic algae play important roles in aquatic primary production, whereas heterotrophic groups feed on bacteria or algae, decompose and cycle organic material or cause diseases in other micro- and macroorganisms, regulating biodiversity and algal blooms^3^. Some protists establish mutualistic associations with corals (e.g., Symbiodinaceae dinoflagellates), hexapods (e.g., Parabasalia and Oxymonadida), other protists (e.g., Foraminifera) and prokaryotes (e.g., methanogenic bacteria and archaea)^4,5^.

Due to genome complexity, cultivation issues and research focus on other microbial groups, the taxonomy and ecology of protists remain poorly understood relative to those of bacteria and fungi^6,7^. Recent advances in environmental DNA (eDNA) analysis, including organelle metagenomics, long-read metabarcoding and single-cell approaches^8–10^ have revealed that the protist world is much more complex than previously thought. Several novel deep lineages, e.g., Hemimastigophora^11^, Provora^12^ and Caelestes^13,14^, have been unearthed, and many more await discovery and formal description^7,15–17^. Omics-based analyses of these novel groups shed light on the early evolution of eukaryotes^14^ and their endosymbioses^10^. Despite the ongoing biodiversity crisis, the vast majority of protist diversity remains undescribed, with major lineages yet to be identified, described and phylogenetically resolved.

This study investigates hitherto undescribed, novel phylogenetic lineages of protists across thousands of soil, sediment and water samples worldwide using long-read eDNA metabarcoding. Our analyses provide phylogenetic placement and DNA-based formal description for 25 phylum-level taxonomic groups, many of which are unexpectedly diverse and widespread.

### Novel protist lineages

Our long-read metabarcoding and phylogenetic analyses revealed 25 coherent, phylum-level lineages of protists, most of which exhibited no affiliations to any currently recognized eukaryote microkingdoms *sensu* Pawlowski^18^ (Figs. 1, S1). Five of these newly discovered phyla formed deeply branching sister lineages within the microkingdoms *-* Koprochomonadia (CRuMs), Haptotellurinophyta (Haptophyta), Parvifortia (Picozoa), Ninturia (Provora) and Aphanistia (Tsukubamonadida). Most of our lineages do not show affiliations to any known kingdoms, suggesting that these lineages in fact may represent kingdoms, although we settle for describing them at the phylum level for now. In other words, our findings nearly double the number of proposed eukaryotic kingdoms or kingdom-level groups, presently at 19-25^9,18–20^, which we successfully reconstructed from SSU-LSU data in most analyses. Since genomes are not available for these novel lineages, we conservatively refer to them at the phylum level in case some of these groups form deep lineages within known kingdoms. These novel phyla mostly branch from the “Excavata” and Holozoa supergroups, but most of them have unstable positions across alternative analyses, consistent with generally weak statistical support for the deepest relationships. Using the DNA-based taxonomy approach^21^, we formally describe 150 novel taxa, including type species in these groups and the corresponding genera, families, orders, classes and phyla (Text S1).

**Fig. 1.**
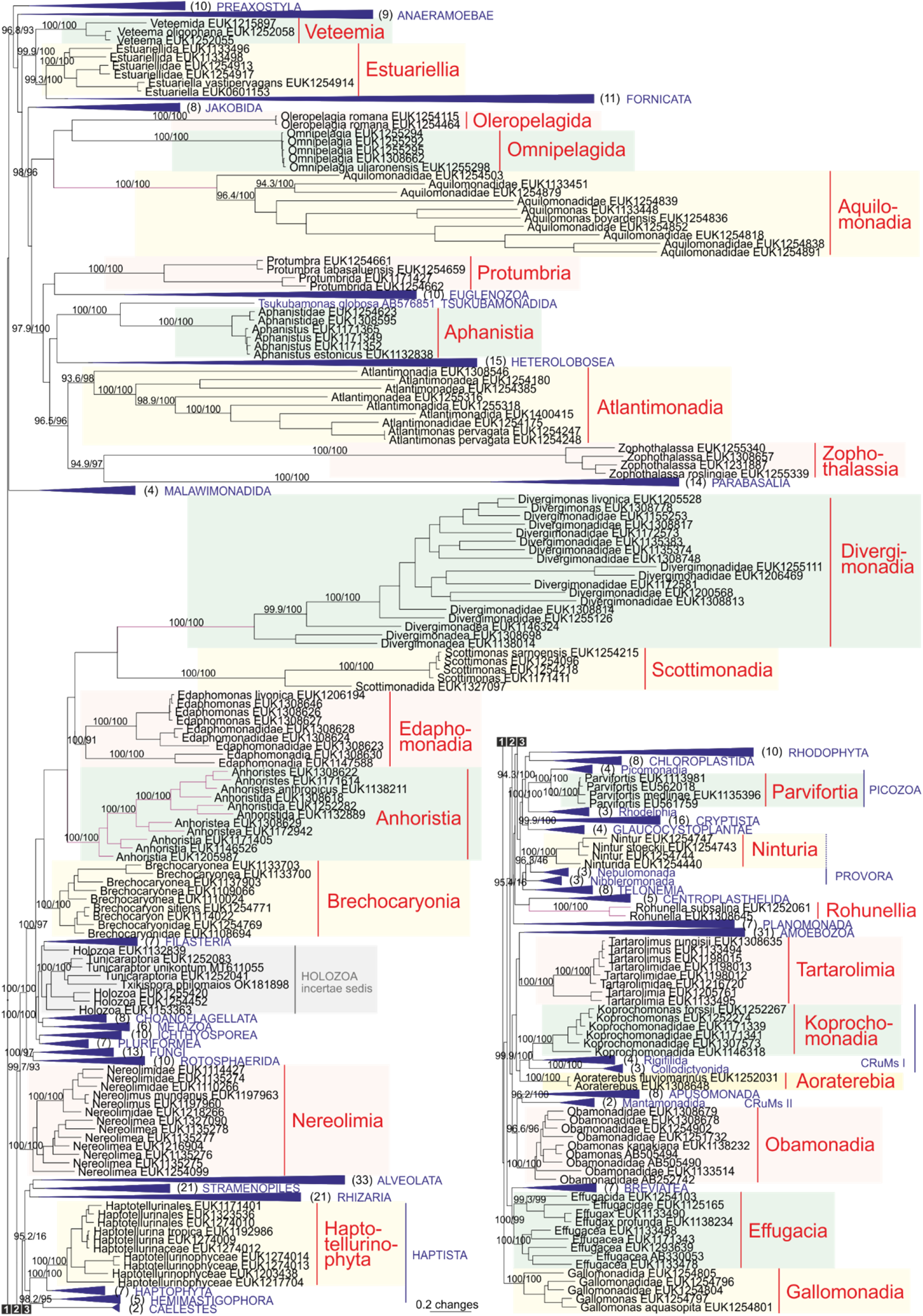
Maximum likelihood tree showing the phylogenetic placement of 25 novel protist phyla (highlighted and named in red typeface) among known eukaryote kingdoms (collapsed branches, names in blue typeface). Representatives of undescribed taxa in Holozoa are highlighted in grey. Numbers in parentheses behind collapsed branches indicate the number of terminal taxa collapsed. The length of branches in purple is reduced by 50%. Numbers above branches indicate rapid bootstrap support in IQ-Tree^24^. All collapsed kingdoms received high support (>95/95); therefore, their support values are not indicated. An uncollapsed tree is provided in Fig. S1.

The predicted species richness of the novel phyla ranges from 1 (Oleropelagida) to 500-760 (Divergimonadia; Data S1). More than 100 species are predicted for Nereolimia (250-380) and Tartarolimia (200-270). Fewer than 10 species are estimated for Aoraterebia (2), Veteemia (3), Parvifortia (4), Omnipelagida (4-6), Protumbria (5) and Scottimonadia (6). In comparison, >100,000 species are described for Archaeplastida (i.e., plants), Fungi and Metazoa, whereas Picozoa and Tsukubamonadida currently harbor a single species. Despite relatively extensive environmental sampling, our approach revealed no representatives of Caelestes. Out of the 25 protist phyla, only representatives of Veteemia were unique to this dataset, whereas 14 phyla had rRNA gene sequence data in INSDC and 20 in the JGI scaffolds database (Data S1; Fig. S2). Compared to <1% chimeric reads in amplicon sequences, 15.0% of the rRNA gene metagenomic scaffolds were chimeric, indicating their limited use in distinguishing biological novelty from analytical artefacts.

Roughly half of our novel phyla are associated with specific substrates, whether or not unequal sampling was accounted for (Fig. 2; Data S2). In particular, soil is the primary habitat (>90% of records) for Anhoristia, Aphanistia, Edaphomonadia, Nereolimia and Tartarolimia, but these phyla show no clear preferences for soil type or land use, suggesting intra-phylum diversity or versatility in terms of vegetation requirements. Interestingly, Haptotellurinophyta, a sister phylum to other groups in photosynthetic Haptophyta, is more common in soil (64% of records) and sediments (26%) than in water. Haptotellurinophyta is probably a different clade from “leptophytes”, a group known based on their plastid and mitochondrial DNA sequences mainly in Arctic waters^10,17^. Several phyla – Aquilomonadia, Brechokaryonia, Oleropelagida, Omnipelagida and Parvifortia – account for >95% of records in marine waters. Furthermore, Aoraterebia occurs across marine, brackish and freshwater systems (92% of records), demonstrating an ability to overcome physiological and ecological barriers posed by salinity. Sediments are the primary habitat for Obamonadia (98% of records in marine sediments) and Estuarellia (91% in freshwater and marine sediments), which may indicate anaerobic or symbiotic (in benthic organisms) lifestyles with potentially amoeboid bodies. Most of the other new phyla are found in soil and water (Atlantimonadia), soil and sediments (Effugacia and Nereolimia), or water and sediments (Gallomonadia and Veteemia). Regarding other substrates, Scottimonadia are common in dung and compost, and Koprochomonadia are common in compost, suggesting a certain level of thermotolerance and potentially host-associated habitat. Protumbria and Divergimonadia occur relatively commonly in decomposing macroalgae, suggesting saprotrophic and potential algal pathogenic interactions. Soil-and sediment-inhabiting groups are geographically widely distributed, but a few small marine groups (Oleropelagida and Omnipelagida) have been recorded only from restricted areas. This indicates either very specific habitat requirements or insufficient sampling of marine environments.

**Fig. 2.**
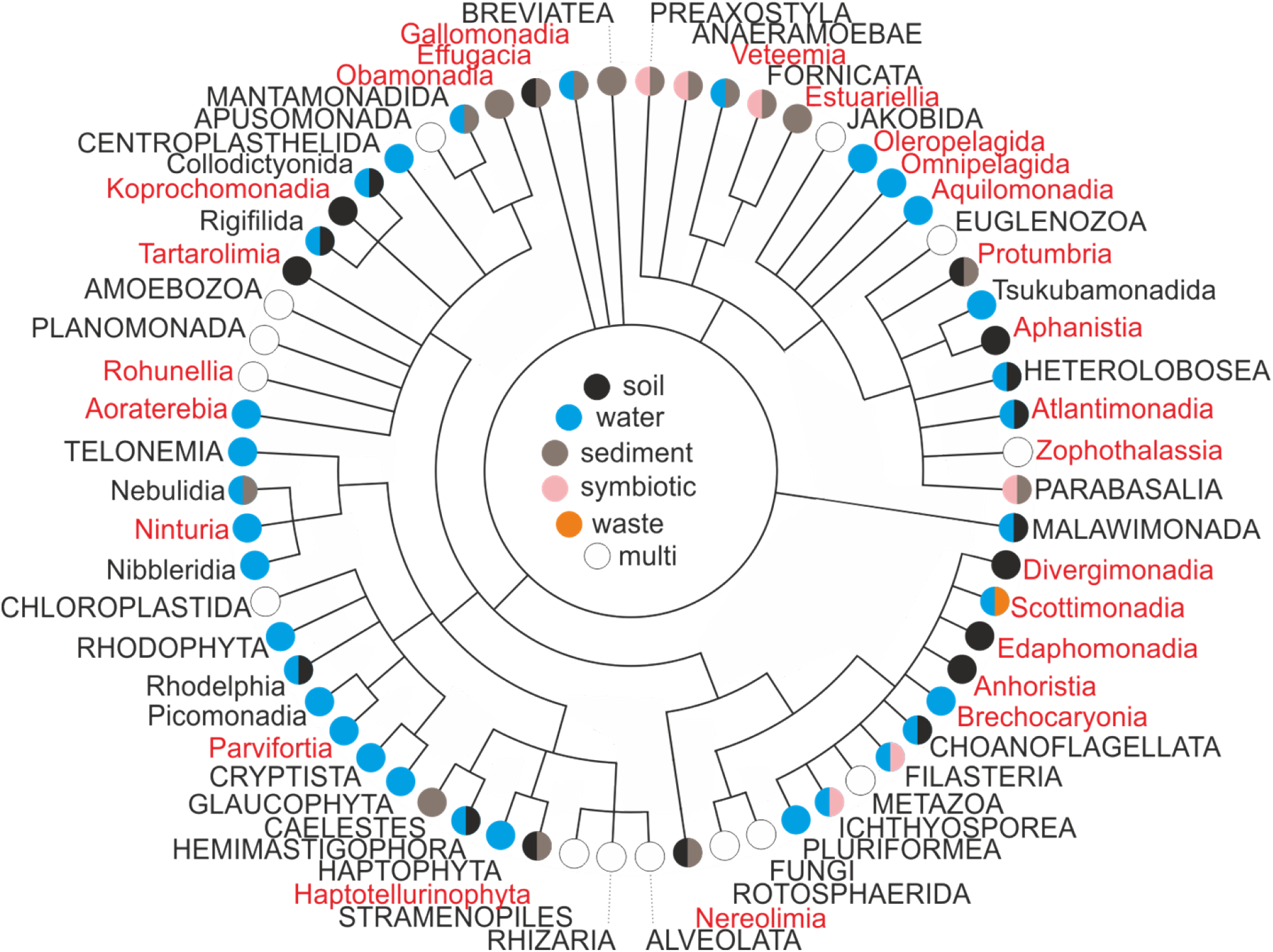
Distribution of novel protist phyla (red) in the eukaryote tree of life, with circles indicating major habitats. Formally recognized microkingdoms are indicated in upper case and phyla in lower case. The figure legend is in the center.

The type species of the novel phyla display great differences in habitat breadth and geographic distribution. *Koprochomonas forssii, Nereolimus mundanus* and *Rohunella subsalina* inhabit multiple substrates, but most type species occur in a single substrate. *Nereolimus mundanus* and *Rohunella subsalina* – but also *Aphanistus estonicus, Nintur stoeckii, Parvifortis medlinae* and *Tartarolimus rungisii* – have a near-global distribution. Nine species are recorded from a single country. Among these, six marine species are documented exclusively in the Marénnes-Oléron basin (Bay of Biscay, France). The high number of unique species may be related to dense local-scale sampling, seasonal population fluctuations or relatively specific environmental conditions, such as oyster farming^22^. Earlier studies similarly identified Sagami Bay in Japan as a hotspot of novel lineages^16^. We assign these lineages to Nereolimia (accession AB505577), Obamonadia (e.g., AB505494) and Gallomonadia (AB505486).

Only 25 novel phyla met the abundance and replication criteria adopted for formal treatment in this study. The largest yet-to-be-described group, “Eukaryota.phy49” (*sensu* EUKARYOME v2.0;^23^), has 132 records from various soil and sediment samples in 45 countries (Data S2). Other such lineages are represented by only one or a few long reads. This limitation is illustrated for a paraphyletic group highlighted as Holozoa *incertae sedis* in Figure 1.

Taken together, our analyses indicate that the number of undescribed phylum- and kingdom-level lineages is at least one-third higher than currently recognized, and many of these are common and diverse components of protist communities in soil, sediments and water, as well as in dung, compost and decomposing macroalgae. We also demonstrate that long reads obtained via metabarcoding can provide an efficient source of phylogenetically informative data for detecting biological novelty. DNA-based taxonomic approaches offer a practical way to recognize and communicate them, giving these taxa the scientific agency their preponderance and ubiquity necessitate. Given the difficulties in culturing many protist groups, the morphoanatomical and functional characterization of these groups will probably remain a matter of high-throughput single-cell microscopic and -omics analyses for at least the next decade.

## Methods

We generated long-read metabarcoding data from eDNA extracts obtained from multiple sources, including 450 soil, 354 sediment and 320 water samples, as well as additional samples from animal dung, compost, decomposing macroalgae and wastewater sludge (Data S3). The DNA of all samples was amplified using the primer pair Euk575F (5’-ascygyggtaaywccagc-3’)^25^ and 21Rngs (5’-gacgaggcatttggctac-3’; shortened from 21R)^26^, designed to cover much of the rRNA cluster and to capture as broad a diversity of eukaryotes as possible while avoiding amplification of prokaryotes. Both primers were tagged with 12-nucleotide indexes for multiplexing^27^. The PCR was performed using TaKaRa Advantage 2 Polymerase Mix (Takara Bio, Kusatsu, Japan), containing 0.5 µl TaKaRa high-fidelity polymerase, 2.5 µl buffer, 0.25 µl dNTP mix (20 mM each), 0.5 µl forward primer (20 µM), 0.5 µl reverse primer (20 µM), 1 µl DNA extract and 19.75 µl ultra-pure water. PCR conditions included an initial activation at 95°C for 2 min, followed by 35 cycles of denaturation at 95 °C for 30 sec, annealing at 68 °C for 4 min, and a final extension at 68 °C for 7 min. PCR products were checked on a 1% agarose-TBE gel electrophoresed at 100 V for 35 min. The DNA samples that did not yield visible amplicons were re-amplified using 38 or 40 cycles. PCR products were pooled based on their intensity on the gel using 1-20 µl of amplicon per sample. Pooled amplicons were cleaned using FavorPrep GEL/PCR Purification Mini Kit (Favorgen Europe, Vienna, Austria) and allocated to 8 PacBio SMRT cell libraries prepared using the SMRTbell Prep kit v3.0 (Pacific Biosciences, Menlo Park, CA, USA), following the manufacturer’s instructions. The libraries were sequenced on 4 25M SMRT cells using the SPRQ Polymerase kit (Pacific Biosciences) and a movie time of 24 hours on the Revio instrument. Circular consensus sequence (CCS) analysis for generating HiFi reads was performed on the instrument. PacBio library preparation and sequencing were performed at the Norwegian Sequencing Centre, Oslo.

PacBio HiFi reads were processed with the NextITS pipeline v1.1.0 (DOI:10.5281/zenodo.15074881)^28^executed in Nextflow v25.10.2^29^. Reads were demultiplexed using LIMA v2.12.0 (Pacific Biosciences) with the ‘--min-score 85’ option. The initial quality filtering discarded sequences containing >4 ambiguous nucleotides, >0.01% expected errors or homopolymer runs longer than 25 nucleotides. Primer trimming was performed with cutadapt v5.1^30^, and reads lacking both primer sites in the correct orientation were removed. The rRNA 18S (SSU), full-length ITS and 28S (LSU) regions were extracted using ITSx v1.1.3^31^ with the ‘all eukaryotes’ setting and using the updated hidden Markov model (HMM) profile database (v2024; profiles are available at https://github.com/Mycology-Microbiology-Center/ITSx_HMMs). The reads were denoised with the UNOISE3 algorithm^32^ with parameters ‘alpha = 6’ and ‘minsize = 1’, and subsequently clustered with VSEARCH at 98% pairwise sequence similarity for the ITS region. The ITS region was selected for clustering because of its higher taxonomic resolution compared to LSU and SSU. Representative ITS sequences were selected based on the greatest similarity to the OTU centroid, then by the mean Phred quality score. Within each sample, sequences differing only in homopolymer length were consolidated, and the most abundant variant was retained as the representative sequence. Within sequencing runs, index-switching (tag-jump) artifacts were filtered with UNCROSS2^33^ using ‘f = 0.01’. To identify OTUs, representative ITS sequences and their corresponding SSU and LSU fragments were separately subjected to BLASTn^34^ searches against the EUKARYOME v1.9.2 database^35^. Identification to kingdom level was considered successful when the best hit had an e-value <e-50 for ITS, or when the best match to a known taxon was >92% in SSU and >90% in LSU. Likewise, identification to the phylum level was considered successful at an e-value <e-60 for ITS, or sequence similarity >94% for SSU and >92% for LSU. These thresholds were intentionally conservative and directed divergent groups within known kingdoms and phyla to additional scrutiny.

Chimera removal was carried out in three stages: (i) *de novo* detection with the UCHIME algorithm^36^ using a maximum chimera score of 0.28; (ii) reference-based validation (uchime_ref) against EUKARYOME; and iii) order-level to kingdom-level consistency in identifications based on SSU, ITS and LSU. When the representative sequence appeared chimeric in SSU or LSU, it was replaced by another ITS representative and the BLASTn-based chimera check was repeated. On several occasions, a chimeric breakpoint was found near the 5’ end of SSU or the 3’ end of LSU, so we ran an additional set of BLASTn searches for the first 300 bases of SSU and the last 1000 bases of LSU. OTUs with a mean Phred score <30 were considered potentially low-quality and removed.

Sequences that could not be confidently matched to the phylum level were included for initial phylogenetic placement analyses. The newly generated reads were supplemented by ∼5000 additional unidentified reads from EUKARYOME, including the long-read dataset of Jamy et al.^1^ and ITS and SSU reads from various sources. As a reference, we downloaded SSU-ITS-LSU or SSU sequences for all known protist genera. Due to the large size of the overall dataset, unidentified sequences were initially aligned and subjected to phylogenetic inference to determine coherent candidate lineages (preliminary numbering available in EUKARYOME v2.0). The sequences were aligned using MAFFT v7.526^37^, followed by manual trimming of overhanging ends, introns, ITS regions and 5.8S rRNA gene and manual correction of apparent misalignment using AliView v1.26^38^. The alignments were processed in ClipKIT v1.4.0^39^ using default options to remove phylogenetically uninformative positions. Phylogenetic analyses were performed using IQ-TREE v2.2.5 (24) with 1000 ultrafast bootstrap replicates. The trees were visualized in FigTree v1.4.4 (*40*). During this process, we eliminated reads that were low-quality (e.g., indels in conserved regions), chimeric or contained long deletions (ca. 15% of all reads) following Tedersoo et al. (*35*). In the second step, we selected up to 10 representative reads per lineage for more detailed phylogenetic analyses using the reference dataset of identified eukaryotes. Roughly two-thirds of the initially unknown lineages were reliably placed to known phyla (mainly Rozellomycota of Fungi, Apicomplexa and Perkinsia of Alveolata, Cercozoa of Rhizaria, Labyrinthulomycota of Stramenopiles and various phyla in Amoebozoa, Euglenozoa and Heterolobosea) and removed from further analyses. The remaining lineages were analyzed iteratively by pruning distant taxa to reduce the dataset size while preserving local phylogenetic context. Finally, we kept only lineages that were well-supported and contained putative species-level taxa represented by at least three reads from different samples, which we considered sufficient for DNA-based taxonomic descriptions. To explore the possible rooting of the eukaryotic tree, we performed additional analyses using various Asgardarchaeota reads as outgroups (*41*), but these produced inconsistent placements and reduced overall branch support. Therefore, we used unrooted trees for the final analyses.

For each of the 25 protist lineages, we prepared separate phylogenetic analyses by compiling all existing sequence data and associated geographical and ecological metadata (as of 30.09.2025), and used *Saccharomyces cerevisiae* (reference strain S298C, accession NR_132207) as an outgroup. When the kingdom-level placement of a lineage was sufficiently clear, we added outgroup taxa from multiple genera within that broader clade. For every eukaryote lineage, we determined the potential type species and assigned terminal taxa into genera, families, orders and classes based on the final phylograms. The type species were selected based on a minimum abundance (>2 reads from different samples), representation by at least one ultra-long read and clear distinction from closely related species. Species diagnoses were prepared based on molecular characters in the SSU, ITS and LSU regions by identifying short, unique diagnostic nucleotide signatures (DNSs) typically 20 bases long, which best distinguished the target species. The DNSs were required to have a minimum number of ambiguous positions for the target species and a maximum number of mismatches to closely related species. Based on the alignments, we estimated the number of allowed differences for the target species to be distinguishable from any related species. Similar DNSs or their combinations were determined for genera, families, orders, classes and phyla.

For the entire alignment length of ITS2 and LSU, we estimated intraspecific sequence variability based on the maximum proportion of differences among individual reads (*21, 23*). To produce rough estimates of potential species richness at the genus and phylum levels, we used manual phylogram assessment and information on ITS intraspecific variability. The clustering threshold for VSEARCH was set to 120% of the type species variability, with a maximum value of 98.0% sequence similarity. We also used the ASAP method (43) implemented in iTaxoTools ASAPy v0.1.2 (*43*) to estimate species-level sequence-similarity thresholds and potential species numbers.

To determine whether the novel phyla had been recovered in previous environmental sequencing efforts, we searched 41,715 publicly available metagenomes and metatranscriptomes from globally distributed environments in the Joint Genome Institute Integrated Microbial Genomes and Microbiomes system, IMG/MER (*44*). For each newly described taxon (N = 167 distinct sequences), we queried the SSU and LSU regions of the rRNA against JGI IMG/MER. For each query sequence and marker, we retained the ten hits with the highest bit scores and then filtered the results to only include alignments longer than 300 bp. In total, 1762 unique reference scaffolds were downloaded. SSU and LSU rRNA regions were identified on these scaffolds using Infernal v1.1.5^45^ with eukaryote-specific covariance model profiles from Rfam (RF01960 for SSU rRNA and RF02543 for LSU rRNA). To detect putative chimeric sequences and refine taxonomic assignments, we performed BLASTn searches for the first and last 500 bp of SSU and LSU against EUKARYOME. Sequences for which different parts matched taxonomically incongruent groups at kingdom and phylum levels were considered chimeric. For phylum-level identification, we applied minimum sequence identity thresholds of 92.0% for SSU and 90.0% for LSU. Only 100% identical matches over at least 500 bases were considered evidence of species-level identity.

The vouchered physical samples that served as a basis for long-read sequence data were selected as holotypes to describe the type species^35,46^. If the physical material had been exhausted during DNA extraction or had otherwise been discarded, the corresponding DNA sample (‘nucleotype’) was nominated instead as holotype^21^. These holotypes were deposited in the repository of the University of Tartu (acronym TUE, with 6-digit accession numbers) unless mentioned otherwise. We also determined ‘legitypes’ (i.e., holotype-derived sequence that is used to represent the species) amongst the longest and highest-quality sequences derived from the nucleotype. Most of the holotypes and underlying environmental samples used for typification originate from composite topsoil samples of the GloSED project^27^ plus water and sediment samples of the ROME^22^ and FunAqua (https://sisu.ut.ee/funaqua/) projects. These samples are linked to the species descriptions and their voucher information in TUE. Legitype and other DNA sequences with metadata were first deposited in the EUKARYOME database (available from v2.0) and subsequently submitted to the UNITE and European Nucleotide Archive databases (accessions OZ474986-OZ475166).

For the naming of taxa, all co-authors were instructed to propose species epithets and generic names independently (Text S2) based on the geographical and ecological metadata and collector information of type material and additional samples. All proposed names were checked for homonymy and grammatical correctness. All co-authors then voted for the most suitable names; in the event of equal votes, the first author decided on the final name, taking into account other names already in use and issued for other lineages.

For establishing higher-ranking taxa, namely genera, families, orders, classes and phyla, we used the following criteria: i) monophyly; ii) bootstrap support >95; iii) phylogenetic breadth and divergence roughly comparable to previously described taxa; and iv) minimizing the number of novel taxa (i.e., preferably retaining larger groups if there were multiple alternative splitting possibilities). Names of higher taxa were deduced from generic names following the International Commission on Zoological Nomenclature^47^ and the International Code of Nomenclature for algae, fungi and plants (the latter for Haptophyta)^48^. The general protist classification follows Adl et al.^19^

## Acknowledgments

We thank Rasmus Puusepp for performing molecular analyses. Sequencing was performed in the Norwegian Sequencing Centre, Oslo. Sequence data retrieved from JGI IMG/MER were produced by the US Department of Energy Joint Genome Institute (https://www.jgi.doe.gov) in collaboration with the user community.

## Author contributions

L.T. and R.H.N designed research; S.S., D.C., M.M., A.M., J.dC., E.L., S.G., K.P., M.B., A.L., M.C.E., R.C., M.H., R.S., M.C., V.P., V.K. and S.A. provided materials and/or data; L.T., V.M., S.S., D.C., K.P., M.M., A.M., J.dC., E.L., A.H., S.G., K.P., M.B., A.M.O., S.R., A.L., V.G.H., M.C.E., R.C., M.H and R.H.N. described and/or named species; V.M., A.H., A.M.O., S.A. and R.M.S. analysed data; L.T. and S.A. provided funding; L.T. and R.H.N. wrote the paper with input from all coauthors.

## Materials and Correspondence

All correspondence and material requests should be addressed to L.T.

## Supplementary Materials

Figs. S1 and S2 Text S1 and S2

References 49–50 (supplement-specific) Data S1-S3

## Data availability

The sequence data are available in EUKARYOME and European Nucleotide Archive. Sequence alignments are available over GitHub. Material and DNA samples are available in TUE. No specific code was written for this analysis.

